# The first poly(A) polymerase from *Alphaproteobacteria*

**DOI:** 10.64898/2025.12.04.692054

**Authors:** Igor P. Oscorbin, Maria S. Kunova, Maxim L. Filipenko

## Abstract

Bacterial poly(A) polymerases (PAPs) play important role in RNA metabolism but remain poorly characterized outside *Gammaproteobacteria*. Here, we cloned and biochemically characterized the first PAP from *Alphaproteobacteria*, specifically from *Marinobacter lipolyticus* (Mli PAP). Using homology-based screening against *E. coli* PAP-1, we identified Mli PAP, sharing 54.8% sequence identity with its *E. coli* counterpart. The enzyme was expressed in *E. coli* but formed insoluble inclusion bodies; active enzyme was solubilized and used for functional assays. Mli PAP exhibited optimal activity at 30°C and similar thermostability to *E. coli* PAP-1. ATP was the preferred substrate, with K_m_ comparable to *E. coli* PAP-1 (1.61 mM and 1.70 mM, respectively), and Mg^2+^ (10 mM) was identified as the optimal cofactor. Mli PAP displayed salt-dependent activity, with the most effective polyadenylation in KCl and inhibition by NaCl and ammonium salts, contrasting with the halophilic nature of its host. This study provides the first functional insights into PAPs from *Alphaproteobacteria*, broadening the understanding of PAP diversity and biochemical properties, potential applications of PAPs in biotechnology.

## 1. Introduction

Polyadenylation in bacterial cells plays a major role in RNA degradation, where polyA tails serve as a signal for RNAses-mediated cleavage. In E. coli, poly(a) talis are mostly synthesized in vivo by a specific poly(A) polymerase (PAP-1), and at least 2 other enzymes are also capable of producing them, including PAP-2 and polynucleotide phosphorylase. Together with closely related CCA-transferases, PAPs are included in the nucleotidyltransferase superfamily, where bacterial PAPs and eukaryotic CCA-transferases stand as class II enzymes, while eukaryotic PAPs and bacterial CCA-transferases are class I enzymes.

Despite early discovery in the 1960s, the details of polyadenylation and its role in cells remain not fully understood [1]. Thus, only 2 bacterial PAPs were cloned and biochemically characterized, namely, PAP-1 from *E. coli* and PAP from *Geobacter sulfurreduc*ens [2–4]. At least 2 other enzymes, *E. coli* PAP-2 [5] and PAP from *Pseudomonas putida* [6], have been also discovered, but these enzymes were purified directly from host organisms without cloning and their functions in cells were not directly studied. Bacterial PAPs were only recovered from *Gammaproteobacteria* (*E. coli* and *P. putida*) and *Desulfuromonadia* (*G. sulfurreducens*). This lack of biochemical data is striking comparing to close counterparts of PAPs, bacterial CCA-transferases that were discovered in a wide range of bacterial taxa: *Gammaproteobacteria, Bacilli, Actinomycetes, Aquificae, Deinococci* and *Cyanophyceae*. While a few CCA-transferases and a few PAP from bacteria were characterized in vitro [7,8], their prediction in silico using amino acid sequences remains a challenging task. Homology of these nucleotidyltransferases is high and only one signature sequence has been reported for bacterial PAPs which does not allow to robustly distinguish PAPs and CCA-transferases [9,10].

The *pcnB* gene encoding bacterial PAPs, is thought to originate from a duplicated CCA-transferase gene and was inherited by *Gamma*- and *Betaproteobacteria* [11]. In some other taxa including *Alphaproteobaceria, Aquificae, Baceroides*, etc, the *pcnB* gene could arise by horizontal transfer, while in other phyla, the gene could be transmitted vertically. Studying putative PAPs outside of *Gammaproteobacteria* can shed light on how these enzymes changed after horizontal transfer. Being a sister class to *Gammaproteobacteria, Alphaproteobacteria* can be a good starting point for these studies because their PAPs are relatively close to *E. coli* PAP-1 and can be identified in silico based on homology to this enzyme. Also, replication of *Alphaproteobacteria* plasmids can also be controlled by the same mechanism involving polyadenylation by PAP as ColE1-type plasmids from *Gammaproteobacteria* [12].

Lately, PAPs became an important tool for biotechnology because of booming development of mRNA vaccines for treatment of various pathological conditions from viral infection to cancer [13,14]. Poly(A) tails can be attached to therapeutic mRNA during transcription being encoded in the respective plasmid. However, this approach is complicated by insertions and deletions in a plasmid sequence encoding poly(A) tail. These altercations lead to length inconsistency of poly(A) tails affecting stability of mRNA. Usage of PAPs allows to overcome this issue but a spectrum of available PAPs is narrow, including yeast PAP and *E. coli* PAP-1. Other bacterial PAP can be superior considering thermal stability and reaction efficacy, yet, these enzymes are not biochemically characterized. Thus, absence of biochemical data not only hampers studying of novel PAPs, but also restricts usage of bacterial PAPs in practical applications.

In the present work, we have cloned and biochemically characterized a novel poly(A)-polymerase from *Marinobacter lipolyticus*, a halophil discovered in saline soil in Spain. This enzyme being the first PAP discovered from *Alphaproteobacteria* was compared with known *E. coli* PAP-1 regarding optimal temperature, thermal stability, salt concentration and cofactor.

## 2 Results

### 2.1 Search for the Mli PAP gene and purification of Mli PAP

To search for new bacterial PAPs, we aligned the amino acid sequence of *E. coli* PAP-1 in Protein BLAST against proteins from *Alphaproteobacteria* using the following criteria: the length 400-600 a.a., the presence of all PAP structural domains (head, neck, body and leg), conservative positions corresponding to D69, D71, E108, characteristic signature of bacterial PAPs [LIV][LIV]G[R/K][R/K]Fx-[LIV]h[HQL][LIV]. Candidate proteins with additional domains were discarded to avoid the presence of other enzymatic activities. The retrieved putative PAPs were ranged based on their similarity to *E. coli* PAP-1, and among 37 candidates, we selected a putative PAP from *Marinobacter lipolyticus* (GenBank: EON91085.1) based on its high similarity to *E. co*li PAP-1 and the host’s habitat. *M. lipolyticus* is an aerobic moderate halophile discovered in saline soil of southern Spain. The optimal growth conditions of 7.5% NaCl, 37°C and pH 7.5 assume a low probability of horizontal gene transfer directly from *E. coli* that growth is inhibited by a 4–7% NaCl. Thus, the gene of *M. lipolyticus* PAP (Mli PAP) has unlikely been transferred from E. coli or its close relatives. Alignment of Mli PAP with other characterized bacterial PAPs: *E. coli* PAP-1 and *Geobacter sulfurreducens* is presented in Figure 1.

**Figure 1.**
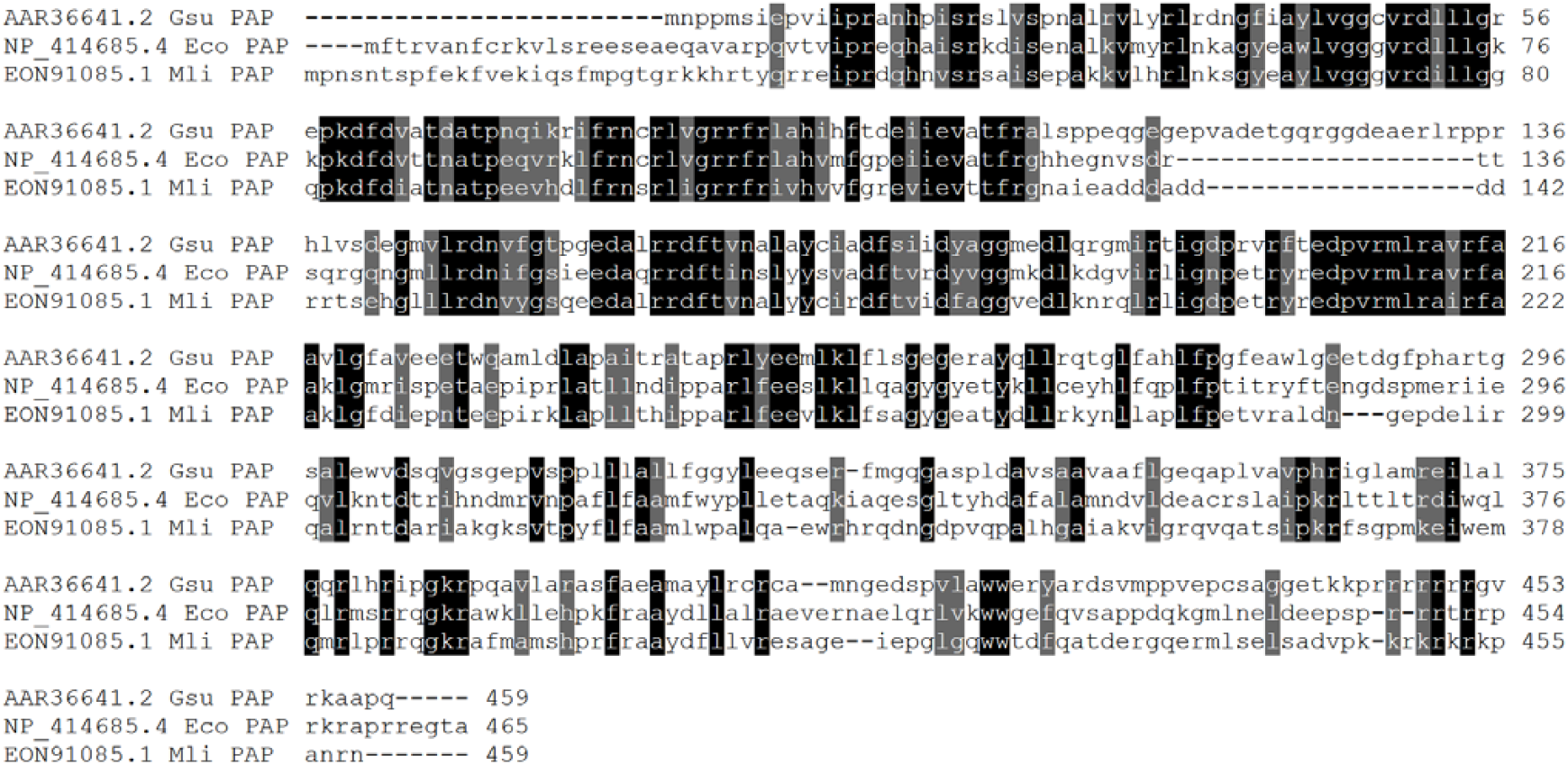
Alignment of Mli PAP and other bacterial PAPs. Amino acid sequences were aligned using ClustalW and visualized in Sequence Manipulation Suite. The colored background indicates conservative amino acid residues.

Homology of the three aligned PAPs is relatively limited: 43.7% between *E. coli PAP-1 and P*AP from *Geobacter sulfurreducens*, 54.8% between *E. coli* PAP-1 and Mli PAP, 43.8% between PAP from *Geobacter sulfurreducens* and Mli PAP. Being one of the search criteria for a new PAP, the [LIV][LIV]G[R/K][R/K]Fx-[LIV]h[HQL][LIV] was conservative among all three PAP. Interestingly, N-terminal half of these proteins was more conservative than the C-terminal part. In *E. coli* PAP-1, C-terminal region is responsible for interplay with degradosome proteins: RNase E, DEAD-box RNA helicases, while N-terminal part hosts catalytically active a.a. residues. Plausibly, the same structure-function relationships can be found in other bacterial PAPs. Also, N-terminus of *Geobacter sulfurreducens* PAP is 20 a.a. shorter than *E. coli* PAP-1 and 20 additional a.a. (positions 115–134) are inserted.

Main properties of the candidate Mli PAP and other bacterial PAPs are listed in Table 1.

**Table 1.**
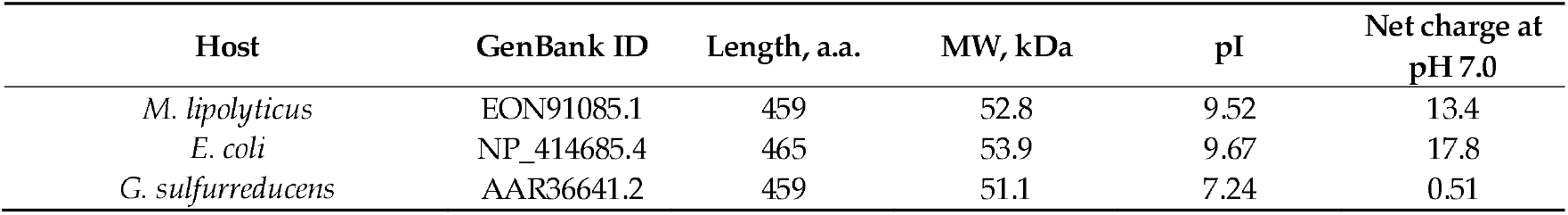
Properties of known bacterial PAPs.

| Host | GenBank ID | Length, a.a. | MW, kDa | pI | Net charge at pH 7.0 |
| --- | --- | --- | --- | --- | --- |
| *M. lipolyticus* | EON91085.1 | 459 | 52.8 | 9.52 | 13.4 |
| *E. coli* | NP_414685.4 | 465 | 53.9 | 9.67 | 17.8 |
| *G. sulfurreducens* | AAR36641.2 | 459 | 51.1 | 7.24 | 0.51 |

While Mli PAP and *E. coli* PAP-1 are highly-charged proteins with a pI value close to 9.5, *G. sulfurreducens* PAP is strikingly different without a high net positive charge Notably, more conservative N-terminal part of the latter protein also has significantly lower net charge than two other PAPs.

After cloning in the pET23a vector, Mli PAP and *E. coli* PAP-1 were expressed in *E. coli BL21* (DE3) pLysS cells following purification of recombinant PAPs. Unfortunately, while *E. coli* PAP-1 purification using a combination of metal-chelate and ion-exchange chromatography was successful, we did not manage to isolate Mli PAP despite all our efforts (Figure 2).

**Figure 2.**
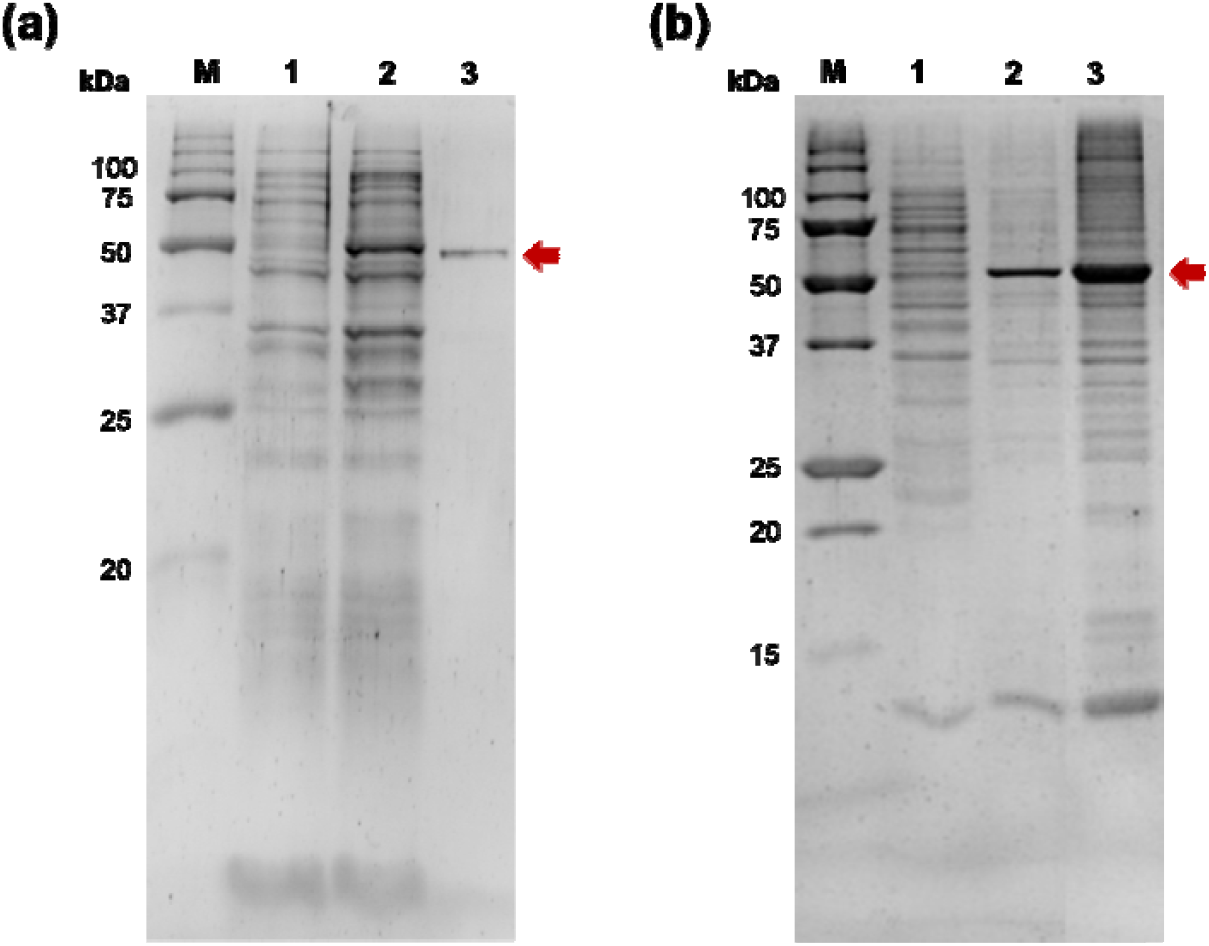
Expression and purification of *E. coli PAP-1 and* Mli PAP. The enzymes were expressed in *E. coli strain BL2*1 (DE3) pLysS. (a) — *E. coli* PAP-1. E. coli PAP-1 was purified using affinity and ion-exchange chromatography. M — Precision Plus Protein standards (Bio-Rad, Hercules, CA, USA), 1 — *E. coli* lysate before expression of *E. coli* PAP-1, 2 — crude lysate after expression, 3 — purified *E. coli PAP-1. (b)* — Mli PAP. Mli PAP was extracted from insoluble inclusion by a high-salt buffer. M — Precision Plus Protein standards (Bio-Rad, Hercules, CA, USA), 1 — soluble protein fraction after expression of Mli PAP, 2 — solubilized Mli PAP, 3 — insoluble inclusion bodies. PAPs are marked by an arrow.

*E. coli* PAP-1 is notoriously known for its tendency to aggregate and being insoluble after lysis, plausibly, because of a high net positive charge. However, a high ionic strength in all purification steps and 2M urea in lysis allowed us to recover active *E. coli* PAP-1 despite its low percent in a soluble protein fraction. The same approach was ineffective for isolation of Mli PAP; virtually all Mli PAP was insoluble after lysis without traceable poly(A)-polymerase activity in supernatant. Similarly to our previous experience with *E. coli* PAP-1, insoluble Mli PAP retained its specific activity allowing to purify the enzyme from inclusion bodies. Up to 20% of active Mli PAP could be extracted from inclusion bodies by resuspension in a high salt buffer. Despite a C-terminal His-Tag, Mli PAP binding with a Ni-charged IMAC resin was ineffective, leaving the most protein in a flowthrow while the eluted Mli PAP was contaminated by cellular proteins to a high degree. Supplying the binding buffer by 2 M urea, 1% Triton X-100, 5 mM beta-mercaptoethanol, or charging IMAC by Co2+ did not allow purification of Mli PAP prisingly, Mli PAP also did not bind with any other used resin, including anion exchanging Macro-Prep HighQ, cation exchanging Macro-Prep HighS, affine Heparin SepFast Media, hydrophobic Macro-Prep t-Butyl HIC in pH 6–9. For that reason, we used solubilized crude Mli PAP with a relatively high amount of contaminant proteins in all further experiments. It should be also noted that no poly(A)-activity was detected in control probes before induction of Mli PAP, ensuring that poly(A)-activity was associated with the recombinant Mli PAP but not host *E. coli* proteins.

### 2.2. Biochemical Properties of Mli PAP

#### 2.2.1. Optimal Temperature and Thermostability

After purification, we studied the biochemical properties of Mli PAP using fluorescently labeled (r)A_20_ oligonucleotide. Here, we assessed the specific activity of Mli PAP in the reaction buffer for *E. coli* PAP-1 and used this estimation for planning further experiments where working conditions assumed less than 5–30% of elongated substrate. Molar activity of Mli PAP was not assessed due to the presence of contaminating proteins.

Reaction temperature is one of the main parameters defining efficacy of amplification. Suboptimal temperature leads to slower poly(A) synthesis or poly(A) polymerase thermal inactivation. To examine the optimal reaction temperature for Mli PAP, we carried out poly(A) tails synthesis at 25–60 °C with a step of 5 °C and quantified the reaction products. The results of the optimal temperature assay are presented in Figure 3.

**Figure 3.**
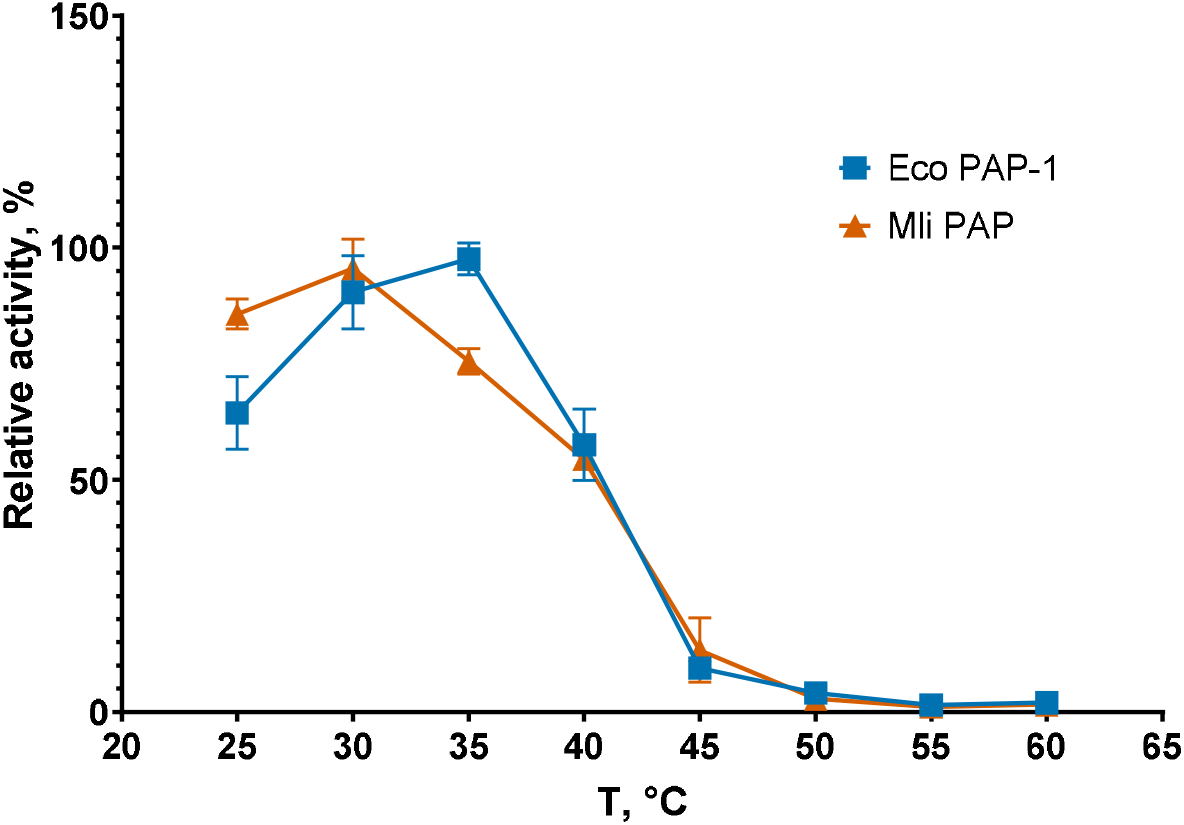
Optimal temperature assay. The optimal temperature for Mli PAP was determined in a poly(A) polymerase activity assay using fluorescently labeled (r)A_20_ oligonucleotide substrate. X-axis marks reaction temperature; Y-axis designates the relative specific activity. Each experiment was triplicated, error bars demonstrate SD.

Optimal temperature was the almost the same for both enzymes, 30°C for Mli PAP and 35°C for *E. coli* PAP-1, which was in match with optimal growth temperature of both host bacteria, 37°C [15]. It should be noted that Mli PAP and *E. coli* PAP quickly lose activity at temperature higher than 35°C becoming almost inactive at 45°C. Next, we tested the thermostability of Mli PAP. Thermal stability defines the ability of an enzyme to remain active after heating. The enzyme was incubated at 30–60 °C for up to 60 min in the reaction buffer for *E. coli* PAP-1 recommended by NEB without ATP following poly(A)-polymerase activity analysis. The results of the assay are presented in Figure 4

**Figure 4.**
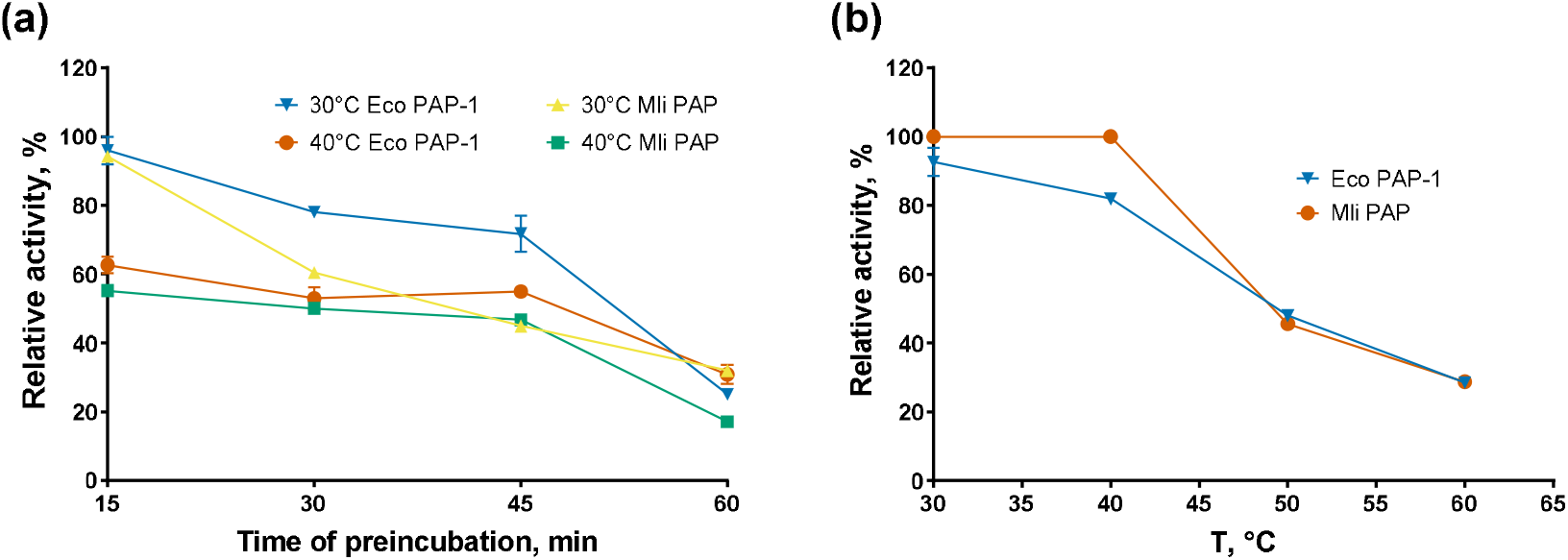
Thermostability assay. (a)—Preincubation without the RNA substrate, (b)— Preincubation with the RNA substrate. Enzymes were incubated at 30–90 °C for up to 60 min following a poly(A)-polymerase activity assay using fluorescently labeled (r)A_20_ oligonucleotide substrate. X-axis marks reaction temperature (a) or the preincubation temperature (b); Y-axis designates the relative specific activity. Each experiment was triplicated, error bars demonstrate SD.

*E. coli* PAP-1 and Mli PAP completely lost activity after incubation for 15 minutes at 50–60 °C and retained only 20–40% of their specific activity after 60 minutes at 30–40 °C, demonstrating relatively low thermal stability. However, both PAPs retained 20–30% of their specific activity in the presence of the (r)A_20_ oligonucleotide substrate after incubation for 60 minutes at 50–60 °C. This stabilizing effect is similar to that observed for reverse transcriptases and DNA polymerases, which are stabilized by RNA and DNA templates, respectively. Thus, prolonged incubation of bacterial PAPs may require supplementation of the incubation buffer with RNA.

#### 2.2.2. Optimal Reaction Buffer

At the next step of Mli PAP characterization, we tested its activity under various conditions including pH, salts, and cofactors. Optimal composition of reaction buffer provides the best possible reaction efficacy, and knowing of enzyme’s ability to synthetize poly(A) tails in varying conditions also allows to perform multiple enzymatic reactions in a single tube. Salt concentration together with pH affect the enzyme’s conformation and its interaction with the respective substrate and other compounds. Cofactors often change specificity of an enzyme enabling chemical transformation of unusual substrates. In our experimental setup, firstly, we titrated Mg^2+^, Mn^2+^, Co^2+^ as cofactors following by titration of salts in various pH. The concentration of the buffer reagent was 50 mM in all the experiments. For divalent cations, the ATP concentration was fixed at 0.5 mM. The results of the cofactor titration experiments are presented in Figure 5.

**Figure 5.**
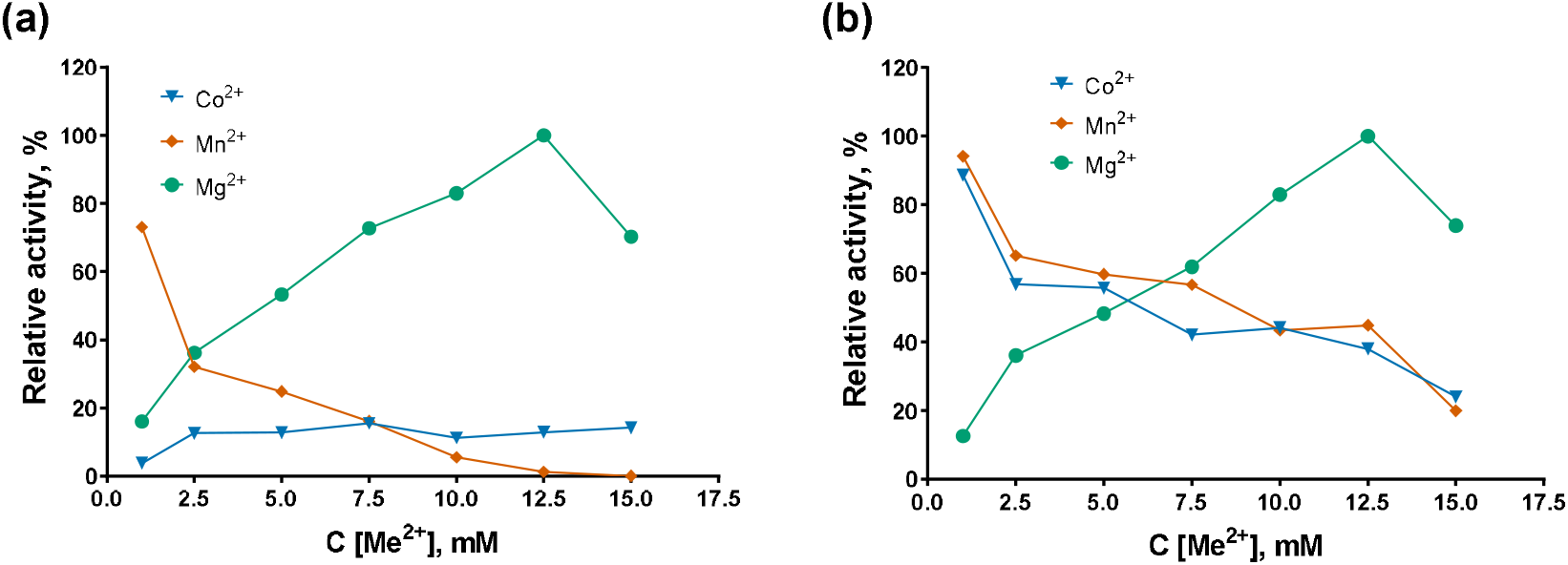
Optimal cofactors. (a)— Mli PAP, (b)— *E. coli* PAP-1. Cofactors were titrated in reaction mixes for a poly(A)-polymerase activity assay using fluorescently labeled (r)A_20_ oligonucleotide substrate. Other components of the reaction buffer were similar to the reaction buffer for *E. coli* PAP-1 recommended by NEB. X-axis marks the cofactors concentrations; Y-axis designates the relative specific activity. Each experiment was triplicated, error bars demonstrate SD.

Among the three tested divalent cations, Mg^2+^ was the preferred cofactor for both enzymes, and the highest activity was observed at a magnesium concentration of 12.5 mM, which was used in all subsequent experiments. *E. coli* PAP-1 was more promiscuous than Mli PAP, efficiently synthesizing poly(A) tails in the presence of 1 mM Co^2+^ or Mn^2+^, retaining 88–95% of its maximal activity. In contrast, Mli PAP retained 73% activity with 1 mM Mn^2+^ but showed less than 20% activity with all tested Co^2+^ concentrations. Plausibly, exchange of magnesium to other cations in PAP’s active site resulted in the decreased activity.

After cofactors, we titrated various salts in the range of 0–200 mM at pH 7.0–9.0 and assessed specific activity of the studied PAPs. The titration results are presented in Figures 6 and 7.

**Figure 6.**
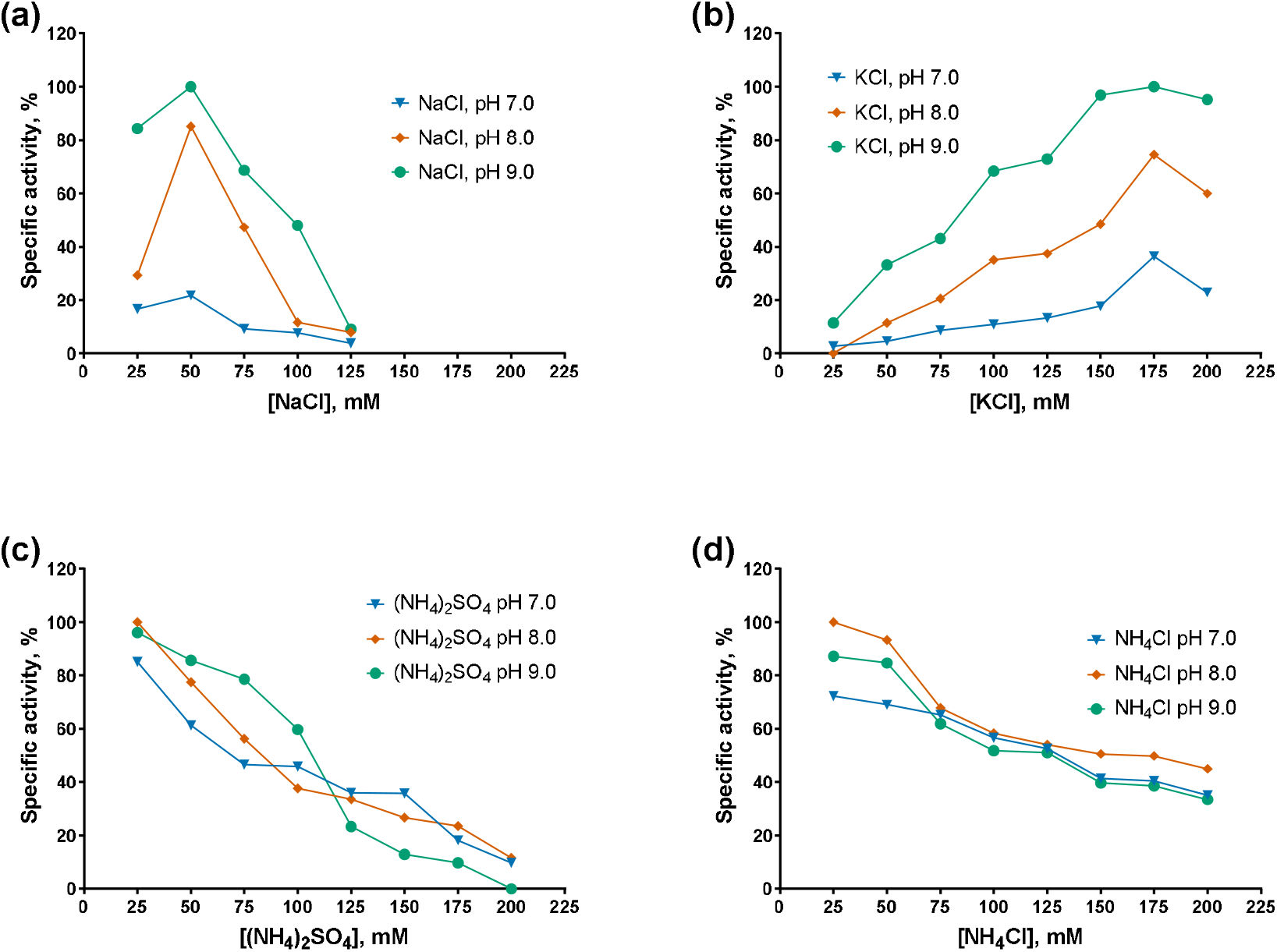
Optimal salts and pH for Mli PAP: (a)—NaCl, (b)— KCl, (c) — (NH_4_)_2_SO_4_, (d) — NH_4_Cl. Salts were salts in the range of 0–200 mM at pH 7.0–9.0 in reaction mixes for a poly(A)-polymerase activity assay using fluorescently labeled (r)A_20_ oligonucleotide substrate. X-axis marks the salts concentrations; Y-axis designates the relative specific activity. Each experiment was triplicated, error bars demonstrate SD.

**Figure 7.**
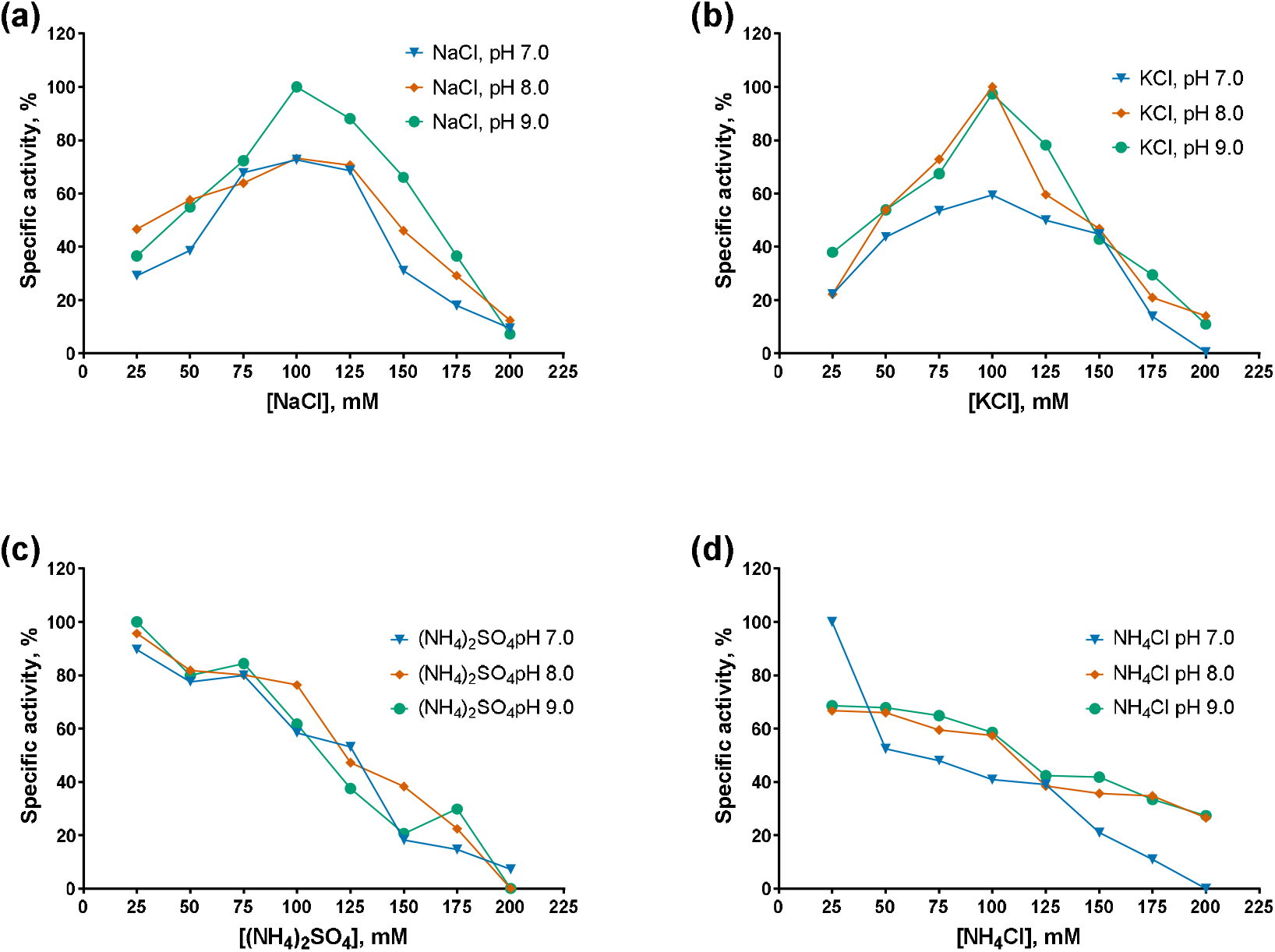
Optimal salts and pH for Mli PAP: (a)—NaCl, (b)— KCl, (c) — (NH_4_)_2_SO_4_, (d) — NH_4_Cl. Salts were titrated in the range of 0–200 mM at pH 7.0–9.0 in reaction mixes for a poly(A)-polymerase activity assay using fluorescently labeled (r)A_20_ oligonucleotide substrate. X-axis marks the salts concentrations; Y-axis designates the relative specific activity. Each experiment was triplicated, error bars demonstrate SD.

The specific activity of both PAPs decreased monotonically with increasing concentrations of ammonium salts, either (NH_4_)_2_SO_4_ or NH_4_Cl, with Mli PAP being slightly more tolerant to NH_4_Cl than *E. coli* PAP-1. In contrast, while *E. coli* PAP-1 exhibited a similar activity profile in the presence of NaCl and KCl (with peak activity at 100 mM salt), Mli PAP lost activity at NaCl concentrations ≥125 mM and showed maximal activity at 150–175 mM KCl. Notably, both enzymes displayed their highest activity in KCl. Regarding pH, both PAPs were the most active at higher pH values in the presence of NaCl and KCl, whereas no clear pH dependence was observed with (NH_4_)_2_SO_4_ or NH_4_Cl. Overall, these results are difficult to reconcile, as inhibition by ammonium ions coincided with the requirement for relatively high KCl concentrations to achieve maximal activity. The preference of Mli PAP for KCl may be related to its tendency to aggregate in low-salt buffers and weak binding to ion-exchange resins. As optimal buffer for Mli PAP we chose 50 mM Tris-HCl, 150 mM KCl, pH 9.0.

#### 2.2.3. Nucleotide Specificity and K_m_ for ATP

Poly(A)-polymerases utilize ATP as the monomer for synthetizing of poly(A) tails. However, *in vitro, E. coli* PAP-1 also inserts CMP, UMP and with a much less efficacy GMP in the growing RNA strand. Thus, *E. coli* PAP-1 is not absolutely specific to ATP and the same could be said for Mli PAP. To test this hypothesis, we titrated all 4 canonical NTP in the range of 0.25–10 mM with a fixed concentration of Mg^2+^ and calculated Km values. The results of the titration experiments are presented in Figure 8.

**Figure 8.**
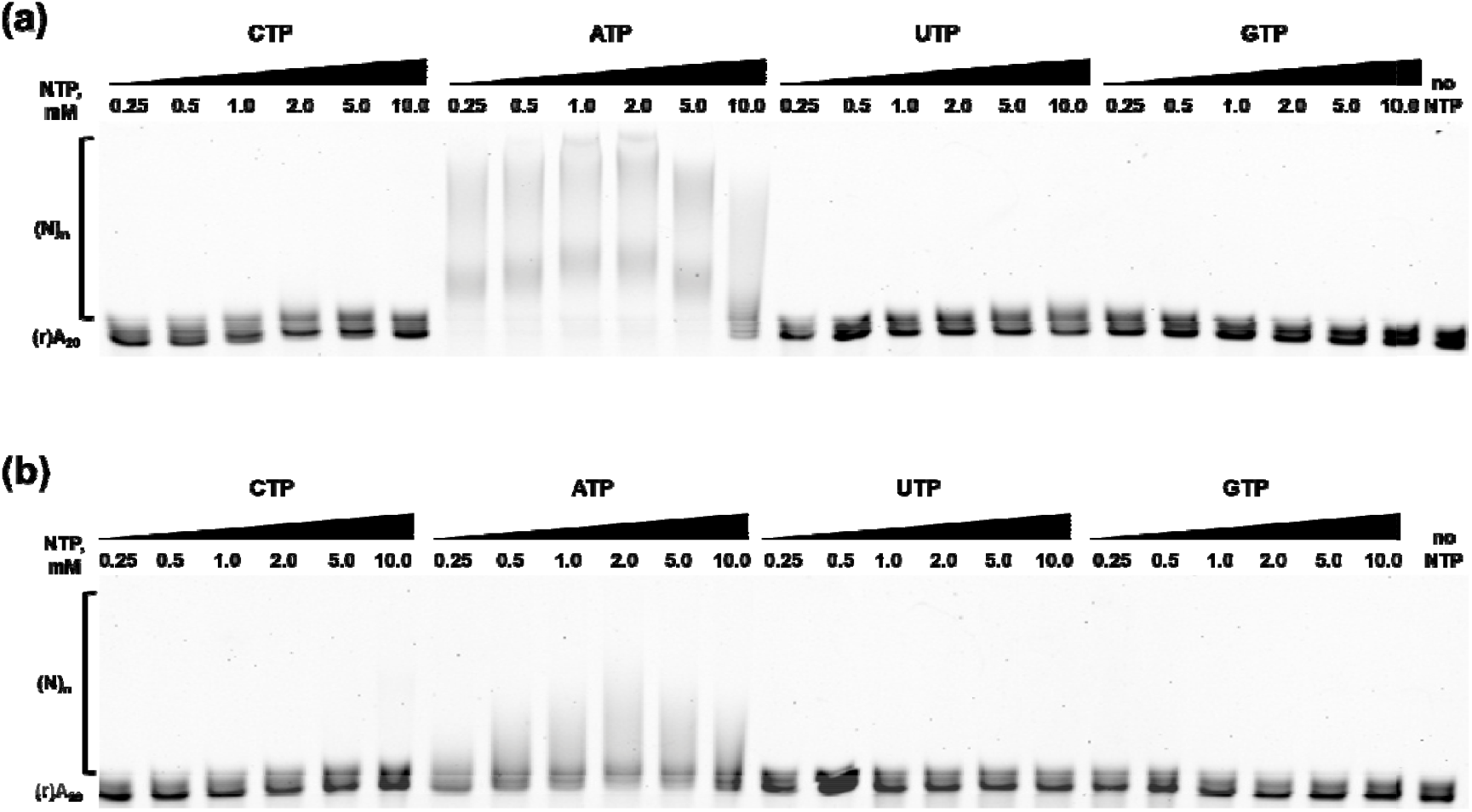
Nucleotide specificity of *E. coli* PAP-1 (a) and Mli PAP (b). NTPs were titrated in the range of 0.25–10 mM in reaction mixes for a poly(A)-polymerase activity assay using fluorescently labeled (r)A_20_ oligonucleotide substrate. NTPs and their concentration are marked above each line, (r)A_20_ demonstrates the substrate’s bands, and (N)_n_ marks the polyadenylated substrate.

As expected, ATP was the preferential substrate for *E. coli* PAP-1 and Mli PAP. Only short elongated products were observed when CTP or UTP were used, while the substrate remained intact with GTP. Saturation was not reached even with 10 mM CTP and UTP demonstrating a discrimination ability of the studied PAPs in our experimental setup, where both enzymes were able to utilize CTP and UTP as a substrate, but with much less efficacy than ATP. Following NTPs titration, we defined K_m_ for ATP by titrating ATP and adjusting reaction time of the polyadenylation assay. The resulted K_m_(ATP) values for *E. coli* PAP-1 and Mli PAP were almost identical—1.70 mM and 1.61 mM—respectively, indicating high degree of similarity between the two enzymes.

## 3. Discussion

Polyadenylation in *E. coli* was discovered in the early 1960s; however, unlike many other enzymes involved in nucleic acid metabolism, PAPs remain understudied. To date, only two bacterial PAPs have been cloned and biochemically characterized, and one additional PAP has been isolated directly from *P. putida*. This scarcity of experimental data makes in silico prediction of new PAPs a non-trivial task, given the high sequence similarity between tRNA nucleotidyltransferases and PAPs. As a result, despite the rapid development of mRNA therapeutics, modern biotechnology currently relies on only two PAPs for in vitro polyadenylation: *E. coli* PAP-1 and yeast PAP. In this study, we selected *Alphaproteobacteria* as a source of new PAPs because members of this class are believed to have acquired PAPs through horizontal gene transfer from *Gammaproteobacteria*, reducing the likelihood of inadvertently cloning a tRNA nucleotidyltransferase instead of a PAP.

*E. coli* PAP is known to be toxic to host cells, and its overexpression can lead to cell death. However, we did not observe any abnormalities in cell growth following induction of either *E. coli* PAP-1 or Mli PAP. It should be noted that we did not specifically assess host-cell viability, and potential toxicity of Mli PAP toward *E. coli* may not manifest as cell lysis. In this context, more detailed studies, including complementation experiments using a pcnB-knockout *E. coli* strain, would be informative and could clarify the impact of a heterologous bacterial PAP on E. coli growth.

Previously, *E. coli* PAP-1 was reported to be prone to aggregation and to require relatively high ionic strength for stability in solution [3,4]. In our experiments, *E. coli* PAP-1 was also largely insoluble after cell lysis and rapidly lost activity when stored in low-salt buffers. A similar effect was observed for Mli PAP, suggesting that poor solubility may be a common characteristic of bacterial PAPs, as their related enzymes, tRNA nucleotidyltransferases, have also been reported to be insoluble [16]. In the study by Shi et al., *E. coli* tRNA nucleotidyltransferase was successfully purified and retained activity after urea denaturation followed by renaturation on Ni–NTA resin. However, neither *E. coli* PAP-1 nor Mli PAP retained activity after a denaturation–renaturation cycle in our hands, so we attempted to purify both enzymes under non-denaturing conditions. A sufficient amount of soluble *E. coli* PAP-1 was available for successful purification. In contrast, Mli PAP expressed in E. coli appeared to be completely insoluble. Fortunately, both PAPs retained detectable activity in inclusion bodies, and we used this observation as the basis for attempts to purify Mli PAP by metal-chelate, ion-exchange, and affinity chromatography. However, Mli PAP solubilized from inclusion bodies using a high-salt buffer failed to bind to any of the tested resins despite extensive variation of pH and ionic strength. Consequently, we characterized the biochemical properties of Mli PAP directly after solubilization from inclusion bodies without further purification, which is one of the main limitations of the present study. It should be emphasized that no poly(A) polymerase activity was detected in *E. coli* cells prior Mli PAP induction, indicating that the results obtained were robust and that no detectable background activity from host poly(A) polymerases interfered with our assays.

Regarding biochemical properties, *E. coli* PAP-1 and Mli PAP demonstrated similar thermostability and optimal temperatures, although their cofactor and salt preferences differed slightly. Both host organisms are mesophiles with optimal growth temperatures near 37 °C; therefore, the observed optimal temperature and thermostability of the PAPs are consistent with expectations for enzymes from mesophilic bacteria. However, *E. coli* PAP-1 showed higher activity in the presence of Mn^2+^ and Co^2+^ and retained more activity in NaCl compared with Mli PAP. Unexpectedly, Mli PAP was strongly inhibited by NaCl but not by KCl, despite its native host, *M. lipolyticus*, tolerating much higher NaCl concentrations than *E. coli* (up to 25% vs. 7–11% w/v, respectively) [15,17]. This discrepancy may hint on stabilization of Mli PAP through interactions with other *M. lipolyticus* proteins that are absent in *E. coli*. The nearly linear inhibition by NH_4_^+^ may relate to the Hofmeister series, in which NH_4_^+^ is among the strongest kosmotropic cations. Given the aggregation tendency of PAPs, inhibition by NH_4_^+^ may reflect salt-induced precipitation. However, this explanation is preliminary and requires direct experimental validation.

Ehe nucleotide specificity of both PAPs was identical: ATP was the preferred substrate, while CMP and UMP were incorporated with much lower efficiency, and GMP incorporation was not detected. This observation partially contradicts previous reports suggesting that *E. coli* PAP-1 can utilize not only ATP but also other NTPs as substrates for RNA elongation [18]. In the study by Yehudai-Resheff and Schuster, the incorporation rates of non-ATP nucleotides were higher than those observed in our experimental system. This discrepancy may be attributed to differences in used reaction buffer compositions and substrates, as we used a short (r)A_20_ oligonucleotide rather than a longer in vitro–transcribed RNA. Nevertheless, both *E. coli* PAP-1 and Mli PAP exhibited relaxed, but not absolute, nucleotide specificity, suggesting the existence of mechanisms that enhance PAP NTPs selectivity in vivo.

Several limitations of the present study should be mentioned. First, the novel PAP from *M. lipolyticus* was isolated with traces of host proteins. Contaminating proteins could potentially influence the observed biochemical properties of Mli PAP and hinder accurate assessment of processivity, molar activity, and turnover number. However, the relatively low sequence similarity between *E. coli* PAP-1 and Mli PAP, together with the absence of detectable residual poly(A) polymerase activity from host cells, makes it unlikely that *E. coli* proteins changed significantly the measured activity of Mli PAP. Second, we did not use long RNA substrates, while RNA secondary structure is known to affect the activity of E. coli PAP-1 [19]. Whether RNA structure modulates poly(A) tail synthesis by Mli PAP remains an open question requiring further investigation. Third, NTP specificity was evaluated qualitatively but not quantitatively, leaving room for speculation regarding subtle differences in substrate preference between the two PAPs. Nevertheless, the similar K_m_(ATP) values measured for *E. coli* PAP-1 and Mli PAP suggest that the both enzymes may incorporate alternative NTPs in a comparable manner.

In summary, we cloned and characterized the biochemical properties of the first poly(A) polymerase identified in *Alphaproteobacteria*, specifically *Marinobacter lipolyticus*. The newly characterized PAP exhibited properties broadly similar to those of *E. coli* PAP-1, providing a foundation for further studies on the diversity and mechanistic features of bacterial PAPs.

## 4. Materials and Methods

### 4.1. Search and selection of Mli PAP coding sequence

Amino acid sequence of *E. coli* PAP-1 was used as a reference for search of potential bacterial PAPs using Protein BLAST against proteins from *Alphaproteobacteria*. Criteria for cloning were length 400-600 a.a., the presence of all PAP structural domains (head, neck, body and leg), conservative positions corresponding to D69, D71, E108 involved in catalysis in PAP I *E. coli* [11], characteristic signature of bacterial PAPs [LIV][LIV]G[R/K][R/K]Fx-[LIV]h[HQL][LIV] [9]. The retrieved putative PAPs were ranged based on their similarity to *E. coli* PAP-1. The selected coding sequence of a putative PAP from *Marinobacter lipolyticus* (GenBank: EON91085.1) was synthesized and cloned into a pET23a vector by Shanghai RealGene Bio-tech, Inc. (Shanghai, China), resulting in a plasmid pPAP-Mli.

### 4.2. Expression and partial purification of PAP from Marinobacter lipolyticus

A starter culture of E. coli BL21 (DE3) pLysS (Promega, WI, Madison, USA) strain harboring the plasmid pPAP-Mli was grown to OD600 = 0.8 in LB medium with 100 μg/mL ampicillin at 37 °C. In a LiFlus GX fermenter (Biotron Inc., Bucheon, South Korea), 4 L of LB with 100 μg/mL ampicillin were inoculated with 20 ml of the starter culture, and the cells were grown to OD600 = 0.6 at 37 °C. The expression of Mli PAP was induced by adding IPTG up to 1 mM concentration. After induction for 12 h at 18 °C, the cells were har-vested by centrifugation at 4,000 × g and stored at −70 °C.

For partial protein purification, the cell pellet was resuspended in the lysis buffer (50 mM Tris-HCl pH 8.0, 300 mM NaCl, 1 mM PMSF, 10 mM DTT, 10% glycerol, 1% Triton X-100, 1 mg/mL lysozyme), incubated for 30 min at 37°C followed by sonication. After lysis, the soluble fraction was separated by two consequent centrifugation steps at 20,000×g for 30 min. The insoluble fraction was resuspended in the resuspension buffer (50 mM Tris-HCl pH 8.0, 1 M NaCl, 1 mM PMSF) and clarified by centrifugation at 25,000×g for 30 min. The resulting supernatant was supplemented by 0.01% NaN_3_ and stored at +4°C. The protein concentration of was measured using a standard Bradford assay.

### 4.3. Expression and partial purification of PAP-1 from E. coli

*E. coli* PAP-1 was cloned into the pET36b vector (Novagen, Madison, WI, USA) using restriction-ligation method and NheI and XhoI restriction sites resulting in the plasmid pET-PAP-Eco as described in our previous work [20]. A starter culture of *E. coli* BL21 (DE3) pLysS harboring the plasmid pET-PAP-Eco was grown to OD600 = 0.6 in LB medium with 100 μg/mL kanamycin at 37 °C. In a LiFlus GX fermenter (Biotron Inc., Bucheon, South Korea), 4 L of LB with 100 μg/mL kanamycin were inoculated with 20 ml of the starter culture, and the cells were grown to OD600 = 0.6 at 37 °C. The expression of *E. coli* PAP-1 was induced by adding IPTG up to 1 mM concentration. After induction for 12 h at 18 °C, the cells were harvested by centrifugation at 4,000 × g and stored at −70 °C.

For protein purification, the cell pellet was resuspended in a lysis buffer (50 mM Tris-HCl pH 8.0, 500 mM NaCl, 1 mM PMSF, 2 M urea, 5 mM β-Mercaptoethanol, 1 mg/mL lysozyme), incubated for 30 min on ice followed by sonication. After lysis, the soluble fraction was separated by two consequent centrifugation steps at 20,000×g for 30 min. Soluble proteins were precipitated ON by 60% (NH_4_)_2_SO_4_, followed by centrifugation at 20,000×g for 30 min. The resulting pellet was suspended in the lysis buffer and loaded onto a 5 ml IMAC column (Bio-Rad, CA, Hercules, USA) pre-equilibrated with a buffer A (50 mM NaPO_4_ pH 7.0, 0.5 M NaCl, 5 mM β-mercaptoethanol), followed by washing the column with 25 ml of the buffer A with 1 M NaCl. Bound proteins were eluted using 10 column volumes and a 0–100% linear gradient of a buffer B (buffer A with 0.5 M imidazole). After affinity chromatography, the fractions with *E. coli* PAP-1 were pooled and loaded onto a 2-ml Macro-Prep HighS Resin (Bio-Rad, column CA, Hercules, USA) pre-equilibrated with a buffer C (50 mM NaPO_4_ pH 7.0, 0.3 M NaCl, 5 mM β-mercaptoethanol). The column was washed with 10 ml of buffer C, and bound proteins were eluted by 10 column volumes and a 0–100% linear gradient of a buffer D (50 mM NaPO_4_ pH 7.0, 1.5 M NaCl, 5 mM β-mercaptoethanol). The fractions with *E. coli* PAP-1 were pooled, dialyzed against a storage buffer (20 mM Tris-HCl, 300 mM NaCl, 0.1 mM EDTA, 1 mM DTT, 0.1% Triton X-100, 50% glycerol, pH 7.5), and stored at −20 °C. All the fractions from each step were analyzed by SDS-PAGE. The purity of the isolated *E. coli* PAP-1 was not less than 95%. The concentration of purified *E. coli* PAP-1 was measured using a standard Bradford assay.

### 4.4. Polyadenylation Assay

The specific activity of poly(A) polymerase was analyzed using elongation of the fluorescently labeled (r)A_20_ oligonucleotide. The reaction mixes (10 μL) contained a 1× reaction buffer for *E. coli* PAP-1 (50 mM Tris-HCl, 250 mM NaCl, 10 mM MgCl_2_, pH 8.0), 1 mM ATP, 10 pmol of (r)A_20_-oligonucleotide, and an indicated amount of PAP. The reactions were started by the addition of the enzyme and immediately transferred to a preheated thermocycler, followed by an incubation for 25 min at 37 °C. After incubation, the reactions were quenched by the addition of 10 μL of formamide and denatured by heating for 5 min at 95 °C. The reaction products were analyzed using denaturing PAGE in a 18% acrylamide gel with 7 M urea and quantified using the ImageLab software (Bio-Rad, Hercules, CA, USA).

### 4.5. Thermal stability, optimal temperature, ion concentration and nucleotide specificity

The thermal stability of Mli PAP and *E. coli* PAP-1 was studied by heating the enzyme and then assaying the poly(A)-polymerase activity. The aliquots of the polymerase activity reaction buffer (described above) without ATP containing an identical amount of Mli PAP were incubated at temperatures from 30°C to 60°C, increasing by 10°C per step for 15–60 min. The reactions were chilled on ice, and the poly(A) polymerase activity was measured as mentioned above.

The temperature optimum of Mli PAP and *E. coli* PAP-1 was defined by measuring the poly(A)-polymerase activity at temperatures from 25°C to 60°C, increasing by 5°C per step, with other conditions identical to those described above for the poly(A) polymerase activity assay. The reactions were initiated by adding the aliquots of enzymes to mixes containing all other components, including the primed template. The mixes were preheated for 5 min before the addition of enzymes.

The optimal ion concentrations were examined as in the poly(A) polymerase activity assay using 25–200 mM NH_4_Cl, KCl, NaCl, (NH_4_)_2_SO_4_ or 1–15 mM MgCl_2_, MnCl_2_, CoCl_2_.

After optimization of reaction conditions, nucleotide specificity of Mli PAP was defined in the optimized reaction buffer (50 mM Tris-HCl, 150 mM KCl, 10 mM MgCl_2_, pH 8.0) by titration of NTPs in reaction mixes for the poly(A) polymerase activity assay in the range of 0.25–10.0 mM. The polyadenylation assay was conducted for 30 min at 37 °C.

### 4.6. Data Analysis

The relative amount of the polyadenylated substrate was calculated as follows—Ratio = I(elongated products)/(I(substrate) + I(elongated products))—and used to estimate by a linear regression model an enzyme amount necessary to elongate 50% of the substrate initially added to the reaction. All calculations were performed in the GraphPad Prism 8.0.1 software (Insight Venture Management, New York, NY, USA).

## 5. Conclusions

In the present study, we successfully cloned and biochemically characterized the first poly(A) polymerase from *Alphaproteobacteria*, specifically, *Marinobacter lipolyticus* (Mli PAP). Despite challenges in solubility and purification, the enzyme retained robust polyadenylation activity when extracted from inclusion bodies. Mli PAP exhibited optimal activity at 30°C, comparable thermostability to *E. coli* PAP-1, strict ATP specificity, and a preference for Mg^2+^ as a cofactor. Notably, it demonstrated distinct salt dependence, with maximal activity in KCl and strong inhibition by NaCl and ammonium salts, highlighting functional divergence from its halophilic host and from *E. coli* PAP-1. These findings expand the biochemical understanding of bacterial PAPs beyond *Gammaproteobacteria* and provide a foundation for exploring PAP diversity, evolutionary mechanisms, and potential biotechnological applications, particularly in RNA-based therapeutics.

## Author Contributions

Conceptualization, M.L.F.; methodology, I.P.O. and M.S.K.; validation, I.P.O.; formal analysis, I.P.O. and M.S.K.; investigation, M.S.K.; resources, M.L.F.; data curation, I.P.O.; writing—original draft preparation, I.P.O.; writing—review and editing, M.L.F.; visualization, M.S.K.; supervision, I.P.O.; project administration, M.L.F.; funding acquisition, I.P.O. All authors have read and agreed to the published version of the manuscript.

## Institutional Review Board Statement

Not applicable.

## Informed Consent Statement

Not applicable.

## Data Availability Statement

Dataset available on request due to the restrictions (e.g., privacy, legal or ethical reasons).

## Abbreviations

PAP: poly(A)-polymerase
Mli: Marinobacter lipolyticus

## Disclaimer/Publisher’s Note

The statements, opinions and data contained in all publications are solely those of the individual author(s) and contributor(s) and not of MDPI and/or the editor(s). MDPI and/or the editor(s) disclaim responsibility for any injury to people or property resulting from any ideas, methods, instructions or products referred to in the content.

